# PandoraGAN: Generating antiviral peptides using Generative Adversarial Network

**DOI:** 10.1101/2021.02.15.431193

**Authors:** Shraddha Surana, Pooja Arora, Divye Singh, Deepti Sahasrabuddhe, Jayaraman Valadi

**Author notes:** Contributing authors.

## Abstract

The continuous increase in pathogenic viruses and the intensive laboratory research emphasizes the need for cost and time efficient drug development. This accelerates research for alternate drug candidates like antiviral peptides(AVP) that have therapeutic and prophylactic potential and gaining attention in recent times. However, diversity in their sequences, limited and non-uniform characterization often limit their applications. Isolating newer peptide backbones with required characteristics is a cumbersome process with many design-test-build cycles. Advanced deep learning approaches such as generative adversarial networks (GAN) can be helpful to expedite the initial stage of developing novel peptide drugs. In this study, we developed PandoraGAN that uses a manually curated training dataset of 130 highly active peptides that includes peptides from known databases (such as AVPdb) and literature to generate novel antiviral peptides. The underlying architecture in PandoraGAN is able to learn a good representation of the implicit properties of antiviral peptides. The generated sequences from PandoraGAN are validated based on physico-chemical properties. They are also compared with the training dataset statistically using Pearson’s correlation and Mann-Whitney U-test. We therefore confirm that PandoraGAN is capable of generating a novel antiviral peptide backbone showing similar properties to that of the known highly active antiviral peptides. This approach exhibits a potential to discover novel patterns of AVP which may have not been seen earlier with traditional methods. To our knowledge this is the first ever use of GAN models for antiviral peptides across the viral spectrum.

## 1 Introduction

The recent viral pandemic has made the entire world fall on its knees. While the efforts for making vaccine were on the forefront, discovery of novel/alternate drug therapies is imperative to keep the disease at bay. Since a few decades, antiviral peptides have emerged to be an attractive candidate against a variety of viruses including RNA viruses like HIV, MERS etc [1]. There have been studies which have proven their prophylactic as well as therapeutic efficiency across some viral classes [2]. Despite numerous studies on antiviral peptides’ isolation, their application in main stream therapeutics is limited due to lack of complete characterization, unclear structure-activity relationship studies, unknown mechanism of action and structural diversity. Use of peptide therapy is often obstructed by their lower stability in the serum and poor availability at the site of infection [2] [3]. Additionally, discovering an effective antiviral peptide candidates against the pathogenic viruses is often a time and labour intensive process. Some dedicated databases like AVPdb [4] and APD3 [5] list the antiviral peptide information along with their efficacy from the literature. However, as wide spectrum of viral hosts diversify, sequence variation is a major hurdle in analysing the existing knowledgebase [6]. Different methods of calculating the potency of these compounds is another drawbacks in comparing the available information.

Therefore there exists an opportunity discover peptides with improved characteristics for potential therapeutic use. Advances in deep learning altered the realm of data driven biological problems and might be helpful to widen existing repertoire of antiviral peptides and to define the specific physicochemical properties and structural patterns relevant to the activity.

GANs have been previously used in biological domain to generate the novel protein sequences from a single family, biological staining images etc. A recent review on the ‘Applications of Generative Adversarial Networks in Drug Design and Discovery’ [7] details the developments using GANs in areas such as drug design and discovery for molecular *de novo* design, dimension reduction task of single-cell data and *de novo* peptide and protein design.

GANs such as the Wasserstein GAN (WGAN) based architecture has been used for generating DNA and tune them to have desired properties [8]. They presented three approaches: generating synthetic DNA sequences using a GAN; a DNA-based variant of the activation maximization design method; and a joint procedure which combines these two approaches together. Another report which focuses on generating synthetic DNA using GAN-based architecture is Feedback-GAN [9] which implements function analyzer to score the generated sequence and pass selected *n* highest scoring sequences to the discriminator’s training dataset as ‘real’ samples. GANs have also been used in the genomics space, for example; using GANs and restricted Boltzmann machines to learn the high dimensional distributions of real genomics datasets and create high quality artificial genomes [10].

Generative models including GANs have proven to be successful in modelling proteins [11]. Protein structures have been generated using GANs, expediting the protein design [12]. A Wasserstein bi-directional Generative Adversarial Networks implemented in HelixGAN, generates structurally plausible solutions of full atom helical structures [13]. GANs have been used to accurately simulate immunogenic peptides with physicochemical properties and immunogenicity predictions similar to that of real antigens [14]. ProteinGAN has been used for generating protein sequences [15]. Their GAN architecture used ResNet blocks in both discriminator and generator model along a single self-attention layer in both. Another report reviews the generative methods and also studies the use of generative methods including GANs for protein sequence design [16]. They depict several studies in *de novo* peptide and protein design using GAN frameworks.

GANDALF which is a GAN-based peptide design system for protein targets uses two networks to generate new peptide sequence and structure [17]. Further research has been carried out with hybrid approach for example, QGAN-HG which is a hybrid GAN with patched quantum circuits, to discover new drug molecules [18]. GANs with adaptive training data have also been used to search for new molecules [19]. Recently, PepGAN [20] has been used to generate antimicrobial peptides showing very high activity.

Research for using GANs and it’s variation and hybrid models for generation of molecules and protein structures is being focused upon. This paper, focuses on using GANs for generating antiviral peptides which form plausible solution and potential starting candidates for further study and engineering of these generated AVPs. PandoraGAN, reported here, was trained using highly active antiviral peptides from the public databases and literature. Additionally, we analysed non-redundant antiviral peptides databases to decide the range of some crucial physico-chemical properties such as hydrophobicity, hydrophilicity, net charge and used these to validate the GAN performance [21, 22]. This study is pioneering in the field of antiviral peptides.

This study can broadly be summarised as:

- Collating anti-viral peptide data
- Training a suitable GAN
- Validating the generated peptides computationally

## 2 Materials & Methods

In this section, we elaborate on the data curation; process of selecting peptides for the training and a set of random non-secretory peptides for a comparative study; and details of PandoraGAN.

### 2.1 Collating anti-viral peptide data

Experimentally validated databases available online like - AVPdb [4], AVPpred benchmark data [23] and peptides in clinical trials were used to build an input dataset. Highly similar sequences that have a similarity greater that 90% were excluded using CDHIT^1^ to get a non redundant dataset. This is to avoid overrepresentation of any particular sequence pattern in the input dataset which may bias GAN output. Sequences with unnatural and modified amino acids such as B (Aspartic acid or Aspargine), J(Leucine or Lsoleucine), O (Pyrrolycine), U (Selenocystine), X (any amino acid) and Z (Glutamic acid or Glutamine)^2^ were also removed as these are out of scope for the current phase of study.

#### 2.1.1 Curation of active and highly active sequences

Antiviral potential is considered to be the key criteria for defining the input dataset.

These are grouped in two separate classes based on their IC50 values [24]. **Active-AVP**: 423 peptide sequences are selected from the databases AVPdb and AVPpred that have an IC50 value less than 30 micro molar as active sequences and refer them as ‘Active-AVP’.

**Highly active sequences (HA-AVP)**: The same data collected from AVPdb is further curated manually based on the peptide’s activity i.e. IC50 values and other information from the literature. Peptides with activity in the range of 0-15 μM are selected to form a subset of highly active peptides. This is an indicative range of highly active peptides as most of the antiviral peptides in clinical trials possess activity in the range of nanomolars. This range would typically represent most of the active peptides from the various class and against various families. The IC50 values are reported in different units in AVPdb ranging from percentage, nano molar, pico molar to qualitative values. Differential reporting of activity values limit the automated curation of a dataset. Hence, the activities are converted to uniform units (micro molar). Peptides from clinical trials as per the recent reports are also added to the dataset.

Subsequently, the sequences with similar identity of more than 90% are removed to ensure equivalent representation of all the peptide patterns. This results in a dataset of 130 peptides which is used for training the PandoraGAN. This set of 130 peptides is referred to as ‘HA-AVP’. The peptides taken from literature and clinical trials can be found in these studies - HIV candidates [25−28]; SARS COV [29]; MERS [30]; T20 [31]; Hepatitis B [32].

#### 2.1.2 Curation of random non-secretory peptides dataset

The dataset consists of 723 random non-secretory peptides with a length of 10- 100 amino acids taken from the Uniprot database [33]. This is used as ‘random peptides’ for comparative study with the PandoraGAN generated peptides.

A summary of the above datasets is listed in Table 1

**Table 1:** A summary of the datasets

| Dataset | Length | IC50 (μM) | #Sequences | Source |
| --- | --- | --- | --- | --- |
| Active-AVP | 7-30 | 0-30 | 423 | AVPdb |
| HA-AVP | 7-30 | 0-15 | 130 | AVPdb and Literature |
| Random non-secretory peptides | 10-100 | NA | 723 | UniProt |

### 2.2 Training a suitable GAN

This section explains the workings of a GAN and further details of LeakGAN which is used in the PandoraGAN architecture.

#### 2.2.1 Generative Adversarial Models

Generative Adversarial Networks (GANs) [34] are deep learning based generative models which learn the underlying data distribution using gradient descent without a requirement of prior knowledge of the structure of the data. With generative and discriminator models working in parallel, GAN iterates between generating the data and testing its equivalence to the input data till the generated data does not appear to be unreal. The GAN model architecture involves two sub-models:

- **Generator model:** It produces samples that mimic the input data distribution. It starts with generating random samples and improves over time by the feedback it receives from the discriminator module.
- **Discriminator model:** Discriminator distinguishes between samples generated by the generator module versus those belonging to the input data. It takes an example (real or generated) as input and predicts a binary class label of real or fake (i.e. generated). The real example comes from the input data on which the GAN is trained and the fake examples are output by the generator model.

The GAN is trained as a two player zero-sum (adversarial) game where the generator model and the discriminator model are considered as the two players from a game theory perspective. In this case, zero-sum means that each of the two players get rewarded when other player does not perform well. This leads to the formulation of minimax loss function on which the GAN is trained and this loss function is defined as

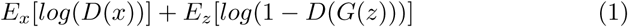

where *G*(.) and *D*(.) are generator and discriminator models respectively, *x* is an input example to the discriminator model and *z* is an input random vector for the generator model.

In order to generate peptides with variable lengths, we use a RNN or LSTM in the generator model - the same concept used for generating variable length textual sentences. Considering an amino acid to be a word and a sequence to be a sentence, the research done in the field of text generation may be modified to generate sequences.

However, for generating text, GAN faces a limitation that the classical discriminative model for the discriminator can only assess a complete sequence. For a partially generated sequence, it is non-trivial to balance the current score and the future score once the entire sequence has been generated. Another problem that is common in text generation using GANs (adversarial modeling) is that the binary signal from the discriminator is not sufficiently informative. This may cause the generator to be inadequately trained and could result in mode collapse problem. Generally, a huge number of training and generated samples are required to improve the generator which becomes very challenging with limited number of experimentally validated AVPs available. To overcome these limitations there are some modified versions of GAN widely used in text generation such as - MaliGAN [35], SeqGAN [36], LeakGAN [37], RankGAN [38], TextGAN [39] and GSGAN [40].

Texygen^3^ which is a text generation model that implements the above mentioned variations of GAN, has been modified to generate antiviral peptides. Experiments were performed with all the above mentioned variations of GAN, and LeakGAN proved to be most successful.

Since the amount of antiviral data we have is very less, the action of leaking the learnt features from the discriminator to the generator is most likely the reason for LeakGAN generating the best set of possible antiviral peptides. LeakGAN is used in PandoraGAN as it helps in deciphering the underlying patterns of antiviral peptides from the limited datasets.

This is described in more detail in the subsection 2.3.

### 2.3 LeakGAN

LeakGAN [37] is a GAN architecture that takes recent advancements in hierarchical reinforcement learning to provide a richer information from the discriminator to the generator. The LeakGAN introduces a hierarchical generator G, which consists of a high-level `Manager` module and a low-level `Worker` module. This helps the architecture to overcome the problem of sparse binary guiding signal which the discriminator provides the generator to learn upon. The `Manager` and `Worker` modules are implemented using Long short-term memory (LSTM) architecture.

In every time step, the `Manager` module receives a high-level feature representation from the discriminator which is used to form the guiding goal for the `Worker` module in that time step. In an adversarial learning, the information within the discriminator and generator models are to be internally maintained. However, in the case of LeakGAN, high-level feature information is passed from discriminator model to generator. This information is said to be the leaked information.

The discriminator is composed of a feature extractor which is a convolutional neural network followed by a dense layer and a sigmoid layer. The information flow in LeakGAN can be seen in Figure 1 where at each step, `Manager` receives the leaked information *f*_*t*_ i.e. the high level features of the input peptides, from the discriminator 𝒟_*φ*_. This is combined with the current hidden state of the `Manager` (i.e. the recurrent hidden vector of the LSTM) to form the goal vector *g*_*t*_. To this goal vector a linear transformation *ψ* is performed to give a *k*-dimensional goal embedding vector 𝒲_*t*_ to incorporate goals produced by the `Manager`.

**Fig. 1:**
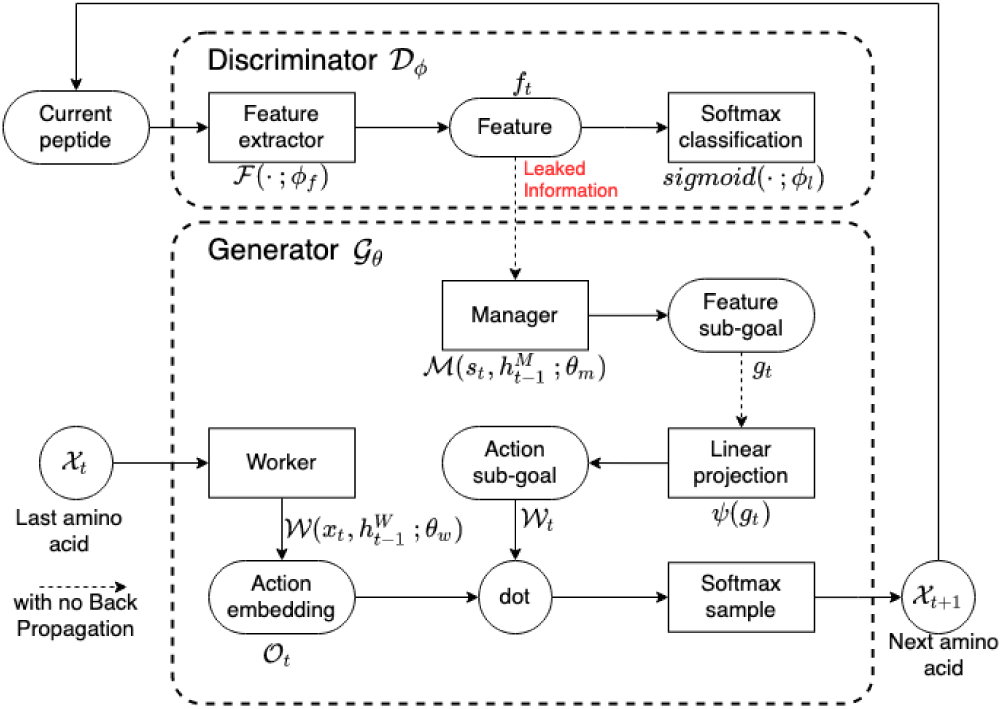
An overview of the LeakGAN framework. The generator is responsible to generate the next amino acid, while the discriminator judges the generated sequence once it is complete. Unlike conventional adverserial training, the discriminator reveals it’s internal state during the training process in order to guide the generator more informatively. Image recreated from the original LeakGAN paper [37]

On the other hand, `Worker` takes the current amino acid residue, *X*_*t*_ as input and outputs a matrix *O*_*t*_ that represents the current vector for all amino acids. This matrix is combined with the goal embedding vector 𝒲_*t*_ to yield the final action space distribution under current state through a softmax.

The LeakGAN is modified to take a sequence of variable length amino acids as input. These form the samples which are termed as ‘real’. The generator generates sequences of random length and amino acid composition which are termed as ‘fake’. The discriminator gets both sets of sequences - the ‘real’ and the ‘fake’ which it will try to correctly classify. With each iteration the discriminator will learn and get more accurate. At the same time, the generator will generate sequences that are more representative of the ‘real’ sequences. For training the model, we use only the sequences themselves. A vocabulary size of 20 is used corresponding to the 20 amino acids.

For generator, to capture the contextual information, single layer LSTM architecture is used for both `Manager` and `Worker`. The `Manager` produces 16- dimensional goal embedding vector 𝒲_*t*_. Further, the input to the GAN gets embedded into a 32-dimensional vector via embedding layer before it goes through the LSTM layer. For discriminator, CNN based architecture is used as feature extractor and classifier. The CNN architecture used in this work has two convolution layers with number of filters in each layers to be 100 and 200 and filter size of 2 and 3, respectively. The convolution layers are followed by a single fully connected dense layer, modeled using highway layer from Highway Network [41]. Finally, a dropout with keep rate of 0.75 and L2 regularisation with lambda value of 0.2 are used to prevent discriminator from overfittig. We pre-train both, the generator and the discriminator modules for 80 epocs. Post this, the adversarial training was done for 100 epocs. The LeakGAN was trained on a machine with 2.2GHz 6-Core Intel Core i7 processor and 2400MHz 32GB DDR4 memory.

The number of peptides to be generated is variable. By default we set this to 100. i.e. once the adversarial model is trained, the generator gives 100 sequences as output. These sequences are given to the post-processing module which filters the sequences based on certain criteria such as - physico-chemical properties viz. net charge at pH 7.4 ≥-1; gravy *>* -1; sequence length between 7 and 30 and secondary structure viz. probability of helix *>* 0.35 and probability of strand *<* 0.3. These threshold have been chosen based on the patterns observed in the input data(HA-AVP). These filtered sequences are the output of PandoraGAN which are further inspected for their composition and patterns.

### 2.4 Validation strategy

The evaluation for artifacts generated from GAN are more qualitative in nature. For e.g. images and text generated via GAN go through human validation. In case of peptides, the approach is to compare distributions with the input training set, or to verify the similarity/overlap with the input training dataset, with the help of PCA. Similar approaches have been used in other papers using GANs [13]. The PandoraGAN generated peptides were compared with the highly active peptides - HA-AVP input training dataset to validate performance of PandoraGAN using the parameters listed in Table 2.

**Table 2:** Validation Strategy Rules that were utilized to validate the peptides generated by PandoraGAN

| Properties | Rules | Values |
| --- | --- | --- |
| Composition | Residues in more than 70% sequences | KR |
|  | Residues in more than 80% sequences | I, L, V |
|  | Residues in less than 20% sequences | M, N |
|  | Residues in around 50% sequences | W, P |
| Physico-chemical | Sequence length | 7 to 30 |
|  | Net charge at Ph7.4 | -1 to 6 |
|  | GRAVY index | > -1 |
| Secondary structure | Probability of Helix | > 0.35 |
|  | Probability of Strand | < 0.3 |
| Repeats and patterns | Dipeptide repeats | FK, KK, LL, VV |
|  | Tripeptide repeats | KKK, KKL, KKV, KKR |
|  | Shannon Entropy of residue | Higher shannon entropy in case of AVP for M, N, P, Y as opposed to non-AVP |
|  | Residue repeats | lower repeats for E, M, N, P as compared to non-AVP |
| Statistical Validation | Correlation of Shannon entropy and residue repeats | high correlation (Pearson's $\rho > 0.9$) |
|  | Mann-Witney U-Test on physio-chemical properties | p-value > 0.05 |

Physico-chemical properties and statistical methods are used to validate the performance of PandoraGAN by comparing it with the HA-AVP set. The validation strategy is shown in Figure 2.

**Fig. 2:**
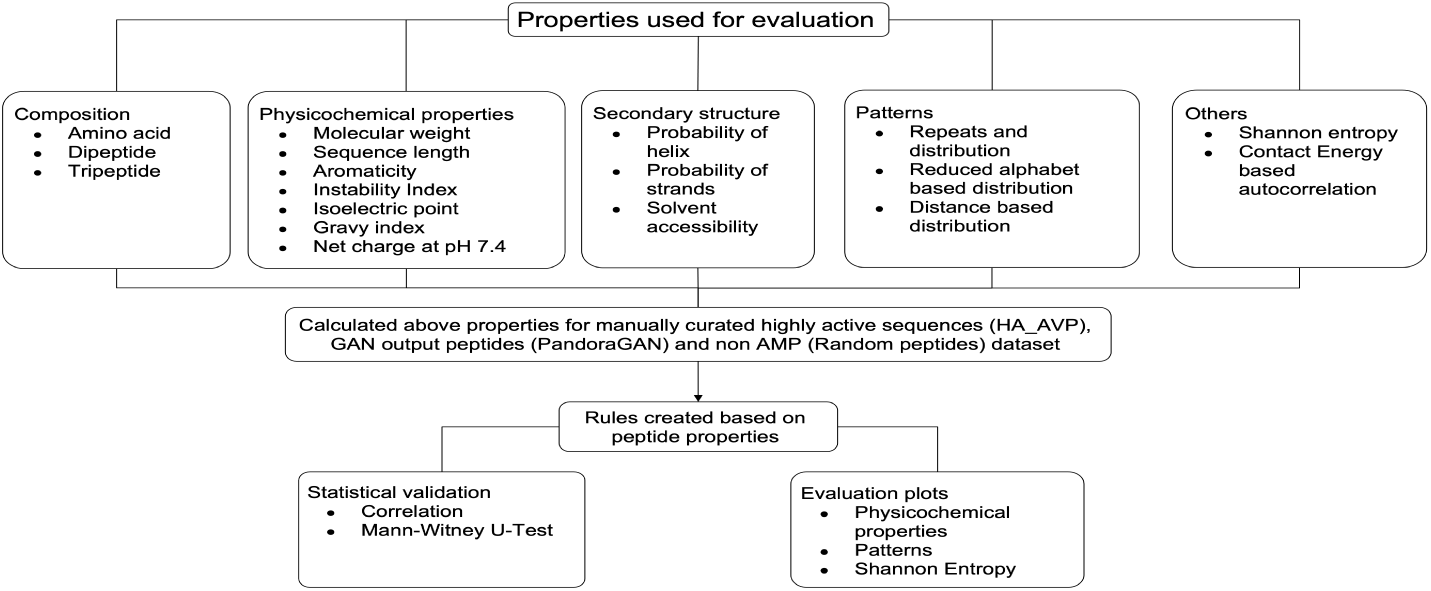
Validation strategy for comparing the peptides using the various properties of highly active peptides.

Physico-chemical properties: Although little elusive, characteristic physico-chemical properties of antiviral peptides have been reported in literature but they lack the crisper boundary conditions for segregation [42]. Various prediction programs use combination of properties like length, charge, hydrophilicity distribution, etc. to classify these peptides. Most of the antiviral peptides are of short length, high in cationic residues and have a spatial distribution of hydrophobic and hydrophillic residues. They also exhibit evolutionary conservation of short motifs which are mostly di, tri or tetra peptide long. These characteristics might be beneficial for exhibiting antiviral activity and reducing the viral load eventually.

To identify the appropriate properties and rules for the dataset extensive analyses of the input training data of highly active peptides (HA-AVP) was performed. Identifiable patterns of such properties were calculated from the input data providing basis for quantitative evaluation criteria. The rules and ranges of physico-chemical properties used in this study are mentioned in Table 2. The parameters utilized in validation strategy were calculated using the python library Biopython [43] and pfeature webserver [44].

Statistical tests: To validate that the physico-chemical properties of peptides generated by PandoraGAN are similar to HA-AVP data, Mann-Whitney U-test (MWU-test) was carried out. MWU-test is a non-parametric equivalent of two sample t-test and has null hypothesis that the two samples come from the same population. As this test is a non-parametric test, it makes no assumption about underlying distributions of the population. Further, to test for similarity in the patterns, pearson’s correlation was calculated for some representative properties.

Figure 3 depicts the overall workflow from dataset creation to validation of the generated peptides. It shows:

**Fig. 3:**
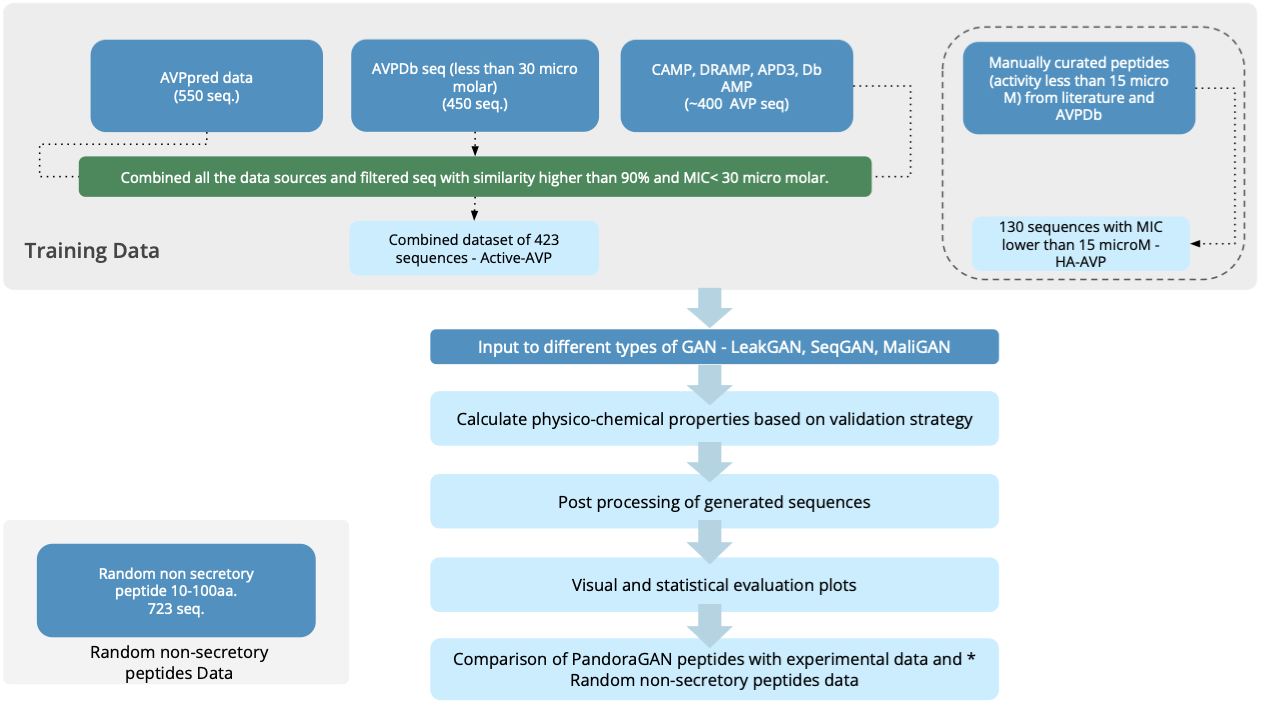
The overall workflow from creating datasets, training and validation of the generated PandoraGAN peptides

- Selection of peptides from various existing databases, creation of the 3 datasets - ‘Active-AVP’; ‘HA-AVP’ and ‘Random peptides’
- Experimentation with different types of GANs by modifying them to take in peptides as inputs
- Calculating the physico-chemical properties and post processing of the generated sequences based on these.
- Creating visual and statistical evaluation plots
- And finally comparing the PandoraGAN generated peptides with the HA- AVP and random peptide dataset.

## 3 Results

PandoraGAN was trained on active-AVP and HA-AVP and the output were compared based on the validation strategy. Here, the determination of activity of peptides and viral hosts is not part of the study. Training the GAN on HA-AVP gives better results. The differentiating factor is the sequences in HA-AVP have lower IC50 values and also contain representation from peptides in clinical trials. Based on the performance of various GAN architecture, we finally implemented LeakGAN to generate novel antiviral peptides in this study. The top five peptides generated by LeakGAN are mentioned in Table 3 sorted by their molecular weight.

**Table 3:** Antiviral peptides sequences generated by PandoraGAN

| S.no | Sequence | Molecular weight | Net charge at pH7.4 |
| --- | --- | --- | --- |
| 1 | ALLEKSKK | 916.12 | 1.60 |
| 2 | VSLVKAALKE | 1057.28 | 0.52 |
| 3 | LLDFKLSDAK | 1149.33 | -0.44 |
| 4 | GAYTFKIRRK | 1239.46 | 3.55 |
| 5 | FKRAKKWVFR | 1365.67 | 4.55 |

The activity data already available was used as an indicator of quality of peptide’s potential and not for determining absolute activity of the generated peptides. In this paper, we report the findings based on the analysis and results of peptides generated by PandoraGAN trained on the HA-AVP dataset and compare it with both the HA-AVP and the random peptides dataset.

The following subsections detail the validation and evaluation criteria based on physico-chemical properties, statistical inferences and cross-validation using existing antiviral classification servers.

### 3.1 Validation based on Physico-chemical properties, repeats and patterns

For evaluating efficiency of PandoraGAN’s learning capabilities, we focus on few physico-chemical properties and compare the generated sequences with the highly active peptides (HA-AVP) as explained in the validation strategy in Figure 2. Various plots utilized to study the comparison of highly active antiviral peptides(HA-AVP), PandoraGAN peptides and random non-secretory peptides are detailed below:

- Amino acid composition: PandoraGAN was able to identify the appropriate composition of amino acids for antiviral peptides. To exemplify our observation, AVP’s are low on Methionine and comparatively high on Lysine and Arginine [21] which is also observed in PandoraGAN generated sequences as seen in Figure 4.

**Fig. 4:**
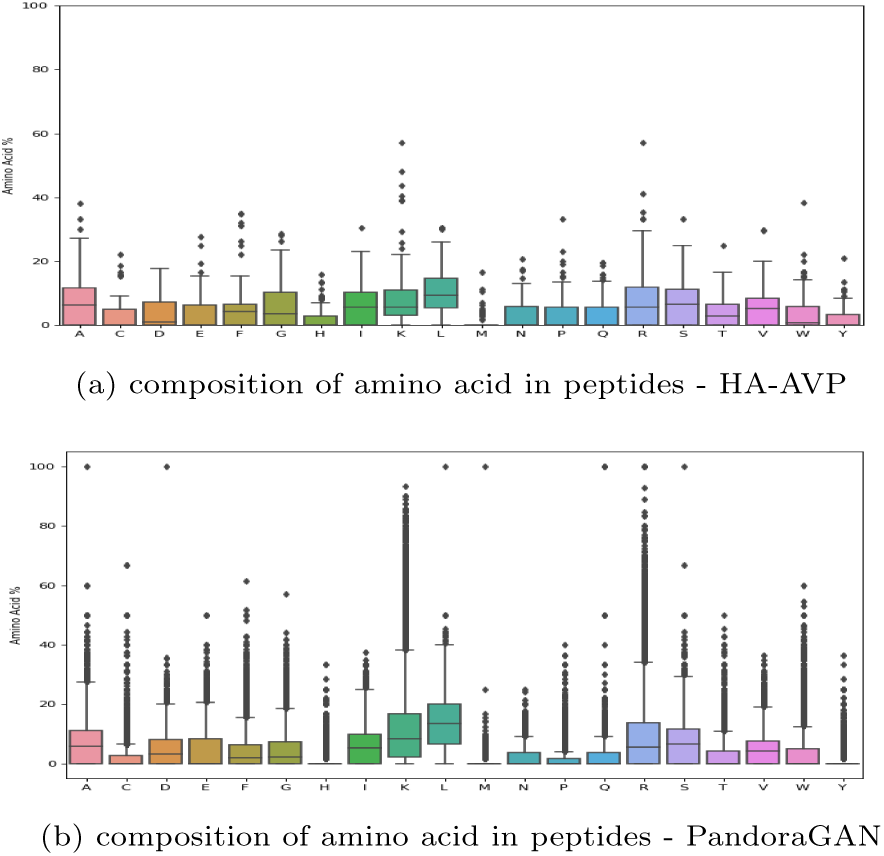
Composition of amino acids in the peptides generated by PandoraGAN. It can be seen that PandoraGAN generated sequences show lower propensity of M, C and high propensity of K,R which is similar to the composition of HA-AVP
- Net charge: PandoraGAN peptides have a comparable range of net charge of -1 to + 4 at pH 7.4, as in the HA-AVP input dataset. This is also validated using MWU-test with p-value=0.116 (*>* 0.05).
- Alpha helix and sheet propensity: Similar to HA-AVPs, the peptides generated by PandoraGAN also possess a higher propensity of alpha helices as compared to sheets which can be explicitly observed in Figure 5a and Figure 5b respectively. It is further validated statistically using MWU-test that the generated peptides have a very similar propensity of alpha helices and sheet sheets with p-values of 0.593 and 0.721 respectively (both *>* 0.05).

**Fig. 5:**
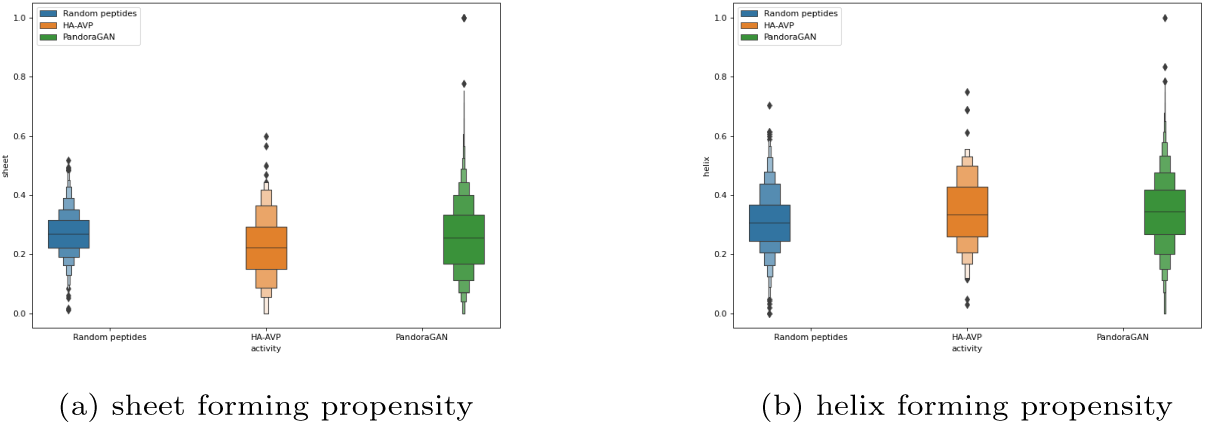
Sequences generated by PandoraGAN and those in HA-AVP dataset follow the same distribution for (a) sheet forming propensity and (b) helix forming propensity as compared to random peptides
- Instability Index: Antiviral peptides are shown to have lower instability index which decreases their chances of in-vivo survival. PandoraGAN generated peptides have an instability index in similar ranges to that of HA-AVP and this is statistically significant with p-value=0.629 (*>* 0.05).
- Shannon entropy and residue repeats: Short amino acid motifs can be crucial for antiviral peptides’ activity. Certain abundant di and tri peptide repeats were identified from the input dataset and compared with Pandora- GAN peptides. The Shannon entropy plots and residue repeats lead to the observation that the entropy of conserved residues in HA-AVP peptides and PandoraGAN peptides is comparable as shown in Figure 6 and Figure 7. The pearson’s correlations between shannon entropy and residue repeats for PandoraGAN peptides and HA-AVP peptides are 0.96 and 0.94 respectively.

**Fig. 6:**
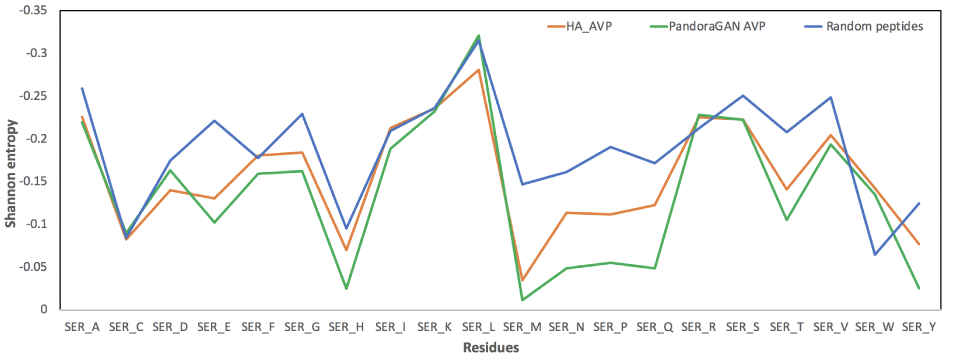
Shannon entropy of all the residues. The Shannon entropy peaks are very similar for PandoraGAN and HA-AVP. It can be seen that both Pandora- GAN and HA-AVP show low peaks for metheonine*ser*_*M*_ which is in abundance in random peptides.

**Fig. 7:**
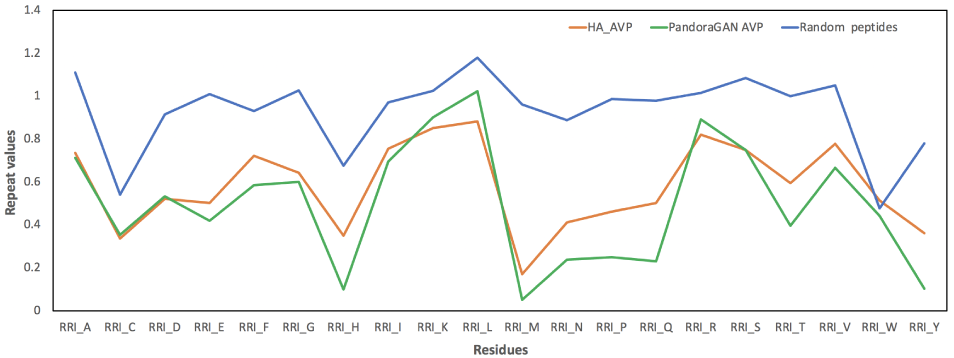
Residue repeats patterns for PandoraGAN generated sequences as compared to HA-AVP sequences. The image shows the residue repeat pattern for random peptides is very different as compared to PandoraGAN generated and HA-AVP set.

These observations confirm the appropriate learning of GAN as the PandoraGAN generated sequences displaying various physico-chemical properties of antiviral peptides, conserved regions and patterns are similar to HA-AVP data.

### 3.2 Statistical Inferences

The comparative analysis of PandoraGAN generated AVPs and the curated HA-AVPs, show that the various properties were highly correlated as can be seen with Pearson’s coefficient mentioned in subsection 3.1. Further, Mann-Witney U-Test shows that aromaticity, instability index, isoelectric point, helix, turn, sheet, gravy and net charge at pH7.4 for PandoraGAN generated AVPs and HA-AVPs follow same distribution with a *p* − *value >* 0.05. These results confirm that the PandoraGAN has learned overall amino acid composition corresponding to physico-chemical properties of HA-AVPs and also validate the peptides generated as output of PandoraGAN. The properties are calculated using the biopython library.

Further, the physico-chemical properties of HA-AVP, PandoraGAN generated sequences and random non-secretory peptides were projected onto first two principal components via principal component analysis. Figure 8 shows an overlap of HA-AVP peptides and PandoraGAN generated peptides while showing clear distinction from random non-secretory peptides.

**Fig. 8:**
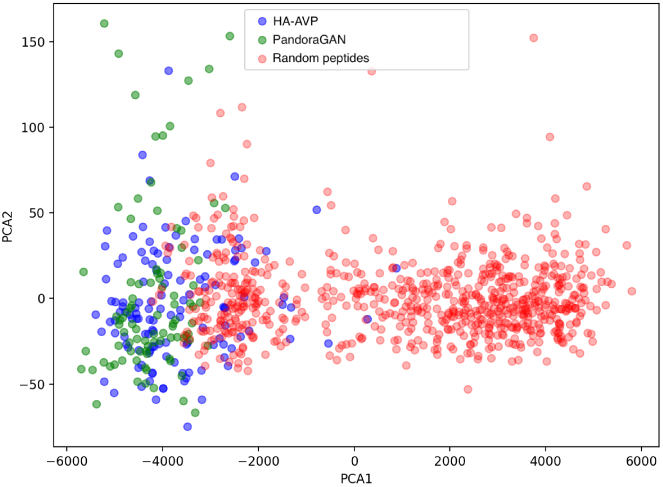
Plot of PCA of the physio-chemical properties of HA-AVP dataset, PandoraGAN generated sequences and random non-secretory peptides. It can be observed that HA-AVP and PandoraGAN generated sequences lie in a similar region in space while the random non-secretory peptides lie in a different region.

### 3.3 Cross-Validation of generated peptides using existing servers

The sequences generated by PandoraGAN and also input data i.e. HA-AVP were also validated by giving them as input to existing antiviral classification servers such as AVPpred [23] and MetaiAVP [45]. It was observed that 70 % of both input data(HA-AVPs) and PandoraGAN generated sequences were, 70% were predicted as antiviral peptides by these servers with a probability greater than 0.9. This additionally validates PandoraGAN’s performance.

## 4 Case study for a specific viral family - Flaviviridae

As a result of diversity in viral families and targets for antivirals, the antiviral peptides also exhibit wide variations in sequence, structure and activities. Sequence based classification techniques many a times can provide limited success in deriving universal features across all the classes of antiviral peptides. Considering this, Pandora GAN was also trained for peptides against specific viral family that may generate more effective antiviral peptides for that particular class and provide the template for wet lab scientists to test effectively. To validate this hypothesis, PandoraGAN was trained on AVPs against flaviviridae family which is one of the well studied class of viruses. Multiple sequences alignment was done with the PandoraGAN output and the input dataset. Out of 10,000 sequences generated by the LeakGAN module, sequences were selected which have molecular weight under 3000 Da, sequence length less than 20 aa and have a higher propensity to form helix (*>*0.3) as shown in flavivirus training dataset. Generated peptides largely contain similar motifs as manifested in the flaviviridae input dataset as observed in Figure 9. Motif patterns like XWLRD are conserved among the input and generated dataset. They also exhibit similar characterization plots as the training data. This result indicates that PandoraGAN outperforms when trained with specific class of antiviral peptides by quickly learning the specific sequence patterns. Therefore we also propose the option of selecting the target viral families for generating effective antiviral peptides using GAN.

**Fig. 9:**
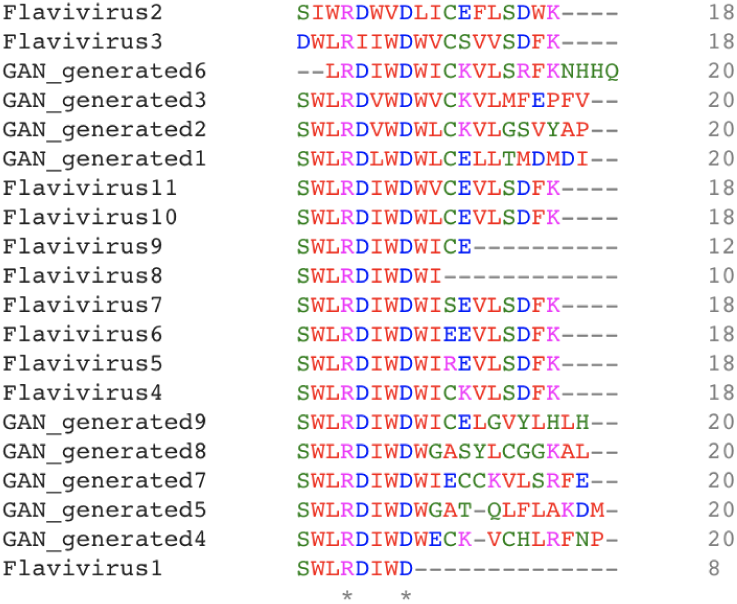
Represents a stretch of Flavivirus sequences where PandoraGAN is able to learn the patterns and create similar sequences.

## 5 Discussion

The dearth in knowledge of antiviral peptides limits potential application of these candidates as alternative therapeutics. Diversity of their host is another concern while generating a specific antiviral peptide. Studying an antiviral peptide in wet-lab setup initiates with identification of viral family, determining the peptide backbone to work on, its source identification, bioinformatics analysis, multiple assays, mutational analysis etc. All these steps make the identification of a novel peptide against specific virus family a laborious and time consuming process. Automation as well as computational advances were only partially successful in easing the load.

PandoraGAN gives a new outlook to the entire process by providing the first step of an effective peptide backbone generated computationally to work on. The generated peptides later can be explored in the laboratory. From our observations, PandoraGAN produces sequences with similar physico-chemical properties to the HA-AVP training data and can be extremely agile depending on the dataset provided. One can also train PandoraGAN using their own training dataset. Computational generation of the peptides would also ensure exploring novel domains of peptides which were otherwise unexplored in the traditional approach.

Here, the physico-chemical properties work as an initial validation for the first line of defense peptides. The amino acid composition, net charge, instability index, repeats and patterns are some of the crucial properties determining the antiviral nature of the peptide.

We also show that PandoraGAN can be used to specifically generate sequences against a particular viral family. Alongwith generating bioactive peptides, another important aspect for the scientist is to know detailed properties of the generated sequences and flexibility to filter and choose the required generated sequence. PandoraGAN provides this ability to the users.

## 6 Conclusion and future work

Here, we present PandoraGAN, a generative model for designing antiviral peptides and demonstrate it’s ability to generate peptides which mimics the distribution of even a small dataset showing prominent characteristics. Generative Adversarial Networks (GAN) based approach for generating antiviral peptides may generate novel motifs for antiviral peptides which could be of therapeutic importance. The method shows that training on a very limited high quality experimental dataset of peptides is capable of producing a robust model that generates peptides which are highly similar to the input training data. In our knowledge, this is the first time GANs have been used for generating antiviral peptides across the viral spectrum.

In future work, we plan to improve PandoraGAN to be able to use both HA-AVP and also the Active-AVP to generate better distribution of residues in a peptide. For this, we will explore techniques that retains the quality of highly active peptides while exposing the GAN to higher number of peptides. We further plan to explore structural and evolutionary aspects of the peptide as reward functions for PandoraGAN to further tune its ability to generate bioactive peptides. This work can further be extrapolated towards identifying the activity of AVPs against specific viral families.

## Authors’ contributions

SS lead the development of the GAN architecture along with Divye S. PA conceptualised the problem and analysed the data. DS provided domain guidance as well as manual dataset curation and JV provided overall guidance.

## Code and Data availability

Antiviral sequences generated by PandoraGAN are available on the PandoraGAN serverhttps://pandora-gan.herokuapp.com/. The code and data is available at https://github.com/thoughtworks/antiviral-peptide-predictions-using-gan/

## Acknowledgments

We would like to acknowledge Govt. of India for launching the Drug Discovery Hacakthon in July 2020 during the COVID 19 to develop computational methods for COVID 19 drug discovery and extrapolate these algorithms to other generic drug discovery challenges.

## Footnotes

1 http://weizhong-lab.ucsd.edu/cdhit-web-server/cgi-bin/index.cgi

2 https://www.ddbj.nig.ac.jp/ddbj/code-e.html

2 https://github.com/geek-ai/Texygen

## References

[1] Vilas Boas, L.C.P., Campos, M.L., Berlanda, R.L.A., de Carvalho Neves, N., Franco, O.L.: Antiviral peptides as promising therapeutic drugs. Cell Mol Life Sci 76(18), 3525–3542 (2019)

[2] Mahendran, A.S.K., Lim, Y.S., Fang, C.-M., Loh, H.-S., Le, C.F.: The potential of antiviral peptides as covid-19 therapeutics. Frontiers in Pharmacology 11, 1475 (2020). https://doi.org/10.3389/fphar.2020.575444

[3] Agarwal, G., Gabrani, R.: Antiviral Peptides: Identification and Validation. Int J Pept Res Ther, 1–20 (2020)

[4] Qureshi, A., Thakur, N., Tandon, H., Kumar, M.: AVPdb: a database of experimentally validated antiviral peptides targeting medically important viruses. Nucleic Acids Res 42(Database issue), 1147–1153 (2014)

[5] Wang, G., Li, X., Wang, Z.: APD3: the antimicrobial peptide database as a tool for research and education. Nucleic Acids Res 44(D1), 1087–1093 (2016)

[6] Di Natale, C., La Manna, S., De Benedictis, I., Brandi, P., Marasco, D.: Perspectives in peptide-based vaccination strategies for syndrome coronavirus 2 pandemic. Frontiers in Pharmacology 11, 1779 (2020). https://doi.org/10.3389/fphar.2020.578382

[7] Lin, E., Lin, C.-H., Lane, H.-Y.: Relevant applications of generative adversarial networks in drug design and discovery: Molecular de novo design, dimensionality reduction, and de novo peptide and protein design. Molecules 25(14) (2020). https://doi.org/10.3390/molecules25143250

[8] Killoran, N., Lee, L.J., Delong, A., Duvenaud, D., Frey, B.J.: Generating and designing DNA with deep generative models. CoRR (2017) 1712.06148

[9] Gupta, A., Zou, J.: Feedback gan (fbgan) for dna: a novel feedback-loop architecture for optimizing protein functions. ArXiv abs/1804.01694 (2018)

[10] Yelmen, B., Decelle, A., Ongaro, L., Marnetto, D., Tallec, C., Montinaro, F., Furtlehner, C., Pagani, L., Jay, F.: Creating artificial human genomes using generative models. BioRxiv (2019). https://doi.org/10.1101/769091

[11] Strokach, A., Kim, P.M.: Deep generative modeling for protein design. Current Opinion in Structural Biology 72, 226–236 (2022). https://doi.org/10.1016/j.sbi.2021.11.008

[12] Anand, N., Huang, P.-S.: Generative modeling for protein structures. In: Proceedings of the 32nd International Conference on Neural Information Processing Systems. NIPS’18, pp. 7505–7516. Curran Associates Inc., Red Hook, NY, USA (2018)

[13] Xie, X., Kim, P.M.: HelixGAN: A bidirectional Generative Adversarial Network with search in latent space for generation under constraints. MLSB (2021)

[14] Li, G., Iyer, B., Prasath, S., Ni, Y., Salomonis, N.: Deepimmuno: deep learning-empowered prediction and generation of immunogenic peptides for t-cell immunity. Briefings in Bioinformatics 22 (2021). https://doi.org/10.1093/bib/bbab160

[15] Repecka, D., Jauniskis, V., Karpus, L., Rembeza, E., Zrimec, J., Poviloniene, S., Rokaitis, I., Laurynenas, A., Abuajwa, W., Savolainen, O., Meskys, R., Engqvist, M.K.M., Zelezniak, A.: Expanding functional protein sequence space using generative adversarial networks. bioRxiv (2019). https://doi.org/10.1101/789719

[16] Wu, Z., Johnston, K.E., Arnold, F.H., Yang, K.K.: Protein sequence design with deep generative models. Current Opinion in Chemical Biology 65, 18–27 (2021). https://doi.org/10.1016/j.cbpa.2021.04.004. Mechanistic Biology * Machine Learning in Chemical Biology

[17] Rossetto, A.M., Zhou, W.: Gandalf: A prototype of a gan-based peptide design method. In: Proceedings of the 10th ACM International Conference on Bioinformatics, Computational Biology and Health Informatics. BCB ‘19, pp. 61–66. Association for Computing Machinery, New York, NY, USA (2019). https://doi.org/10.1145/3307339.3342183. https://doi.org/10.1145/3307339.3342183

[18] Li, J., Topaloglu, R.O., Ghosh, S.: Quantum generative models for small molecule drug discovery. IEEE Transactions on Quantum Engineering 2, 1–8 (2021). https://doi.org/10.1109/TQE.2021.3104804

[19] Blanchard, A.E., Stanley, C., Bhowmik, D.: Using gans with adaptive training data to search for new molecules. J Cheminform 13 (14) (2021). https://doi.org/10.1186/s13321-021-00494-3

[20] Tucs, A., Tran, D.P., Yumoto, A., Ito, Y., Uzawa, T., Tsuda, K.: Generating Ampicillin-Level Antimicrobial Peptides with Activity-Aware Generative Adversarial Networks. ChemRxiv (2020). https://doi.org/10.26434/chemrxiv.12116136.v1

[21] Chang, K.Y., Yang, J.-R.: Analysis and prediction of highly effective antiviral peptides based on random forests. PloS one 8(8), 70166 (2013)

[22] Beltrán Lissabet, J.F., Belén, L.H., Farias, J.G.: AntiVPP 1.0: A portable tool for prediction of antiviral peptides. Comput Biol Med 107, 127–130 (2019)

[23] Thakur, N., Qureshi, A., Kumar, M.: AVPpred: collection and prediction of highly effective antiviral peptides. Nucleic Acids Res 40(Web Server issue), 199–204 (2012)

[24] Qureshi, A., Tandon, H., Kumar, M.: AVP-IC50 Pred: Multiple machine learning techniques-based prediction of peptide antiviral activity in terms of half maximal inhibitory concentration (IC50). Biopolymers 104(6), 753–763 (2015)

[25] Liang, X., Zhang, X., Lian, K., Tian, X., Zhang, M., Wang, S., Chen, C., Nie, C., Pan, Y., Han, F., Wei, Z., Zhang, W.: Antiviral effects of Bovine antimicrobial peptide against TGEV in vivo and in vitro. J Vet Sci 21(5), 80 (2020)

[26] Shi, S., Nguyen, P.K., Cabral, H.J., Diez-Barroso, R., Derry, P.J., Kanahara, S.M., Kumar, V.A.: Development of peptide inhibitors of hiv transmission. Bioactive materials 1(2), 109–121 (2016)

[27] Chupradit, K., Moonmuang, S., Nangola, S., Kitidee, K., Yasamut, U., Mougel, M., Tayapiwatana, C.: Current peptide and protein candidates challenging hiv therapy beyond the vaccine era. Viruses 9(10) (2017). https://doi.org/10.3390/v9100281

[28] Coffey, M.J., Woffendin, C., Phare, S.M., Strieter, R.M., Markovitz, D.M.: RANTES inhibits HIV-1 replication in human peripheral blood monocytes and alveolar macrophages. Am J Physiol 272(5 Pt 1), 1025–1029 (1997)

[29] Xia, S., Liu, M., Wang, C., Xu, W., Lan, Q., Feng, S., Qi, F., Bao, L., Du, L., Liu, S., et al.: Inhibition of sars-cov-2 (previously 2019-ncov) infection by a highly potent pan-coronavirus fusion inhibitor targeting its spike protein that harbors a high capacity to mediate membrane fusion. Cell research 30(4), 343–355 (2020)

[30] Zhao, H., Zhou, J., Zhang, K., Chu, H., Liu, D., Poon, V.K., Chan, C.C., Leung, H.C., Fai, N., Lin, Y.P., Zhang, A.J., Jin, D.Y., Yuen, K.Y., Zheng, B.J.: A novel peptide with potent and broad-spectrum antiviral activities against multiple respiratory viruses. Sci Rep 6, 22008 (2016)

[31] Ding, X., Zhang, X., Chong, H., Zhu, Y., Wei, H., Wu, X., He, J., Wang, X., He, Y.: Enfuvirtide (T20)-Based Lipopeptide Is a Potent HIV-1 Cell Fusion Inhibitor: Implications for Viral Entry and Inhibition. J Virol 91(18) (2017)

[32] Lempp, F.A., Qu, B., Wang, Y.-X., Urban, S.: Hepatitis b virus infection of a mouse hepatic cell line reconstituted with human sodium taurocholate cotransporting polypeptide. Journal of virology 90(9), 4827–4831 (2016)

[33] Consortium, T.U.: UniProt: a worldwide hub of protein knowledge. Nucleic Acids Research 47(D1), 506–515 (2018) https://academic.oup.com/nar/article-pdf/47/D1/D506/27437297/gky1049.pdf. https://doi.org/10.1093/nar/gky1049

[34] Goodfellow, I.J., Pouget-Abadie, J., Mirza, M., Xu, B., Warde-Farley, D., Ozair, S., Courville, A., Bengio, Y.: Generative adversarial nets. In: Proceedings of the 27th International Conference on Neural Information Processing Systems - Volume 2. NIPS’14, pp. 2672–2680. MIT Press, Cambridge, MA, USA (2014)

[35] Che, T., Li, Y., Zhang, R., Hjelm, R.D., Li, W., Song, Y., Bengio, Y.: Maximum-likelihood augmented discrete generative adversarial networks. CoRR (2017) 1702.07983

[36] Yu, L., Zhang, W., Wang, J., Yu, Y.: Seqgan: Sequence generative adversarial nets with policy gradient. CoRR (2017) 1609.05473 [cs.LG]

[37] Guo, J., Lu, S., Cai, H., Zhang, W., Yu, Y., Wang, J.: Long text generation via adversarial training with leaked information. CoRR (2017) 1709.08624

[38] Lin, K., Li, D., He, X., Zhang, Z., Sun, M.-t.: Adversarial ranking for language generation. In:Guyon, I., Luxburg, U.V., Bengio, S., Wallach, H., Fergus, R., Vishwanathan, S., Garnett, R. (eds.) Advances in Neural Information Processing Systems, vol. 30, pp. 3155–3165. Curran Associates, Inc., ??? (2017). https://proceedings.neurips.cc/paper/2017/file/bf201d5407a6509fa536afc4b380577e-Paper.pdf

[39] Zhang, Y., Gan, Z., Fan, K., Chen, Z., Henao, R., Shen, D., Carin, L.: Adversarial feature matching for text generation. In: International Conference on Machine Learning, pp. 4006–4015 (2017). PMLR

[40] Kusner, M.J., Hernández-Lobato, J.M.: Gans for sequences of discrete elements with the gumbel-softmax distribution. arXiv preprint 1611.04051 (2016)

[41] Srivastava, R.K., Greff, K., Schmidhuber, J.: Highway networks. CoRR (2015) 1505.00387 [cs.LG]

[42] Kang, S.J., Kim, D.H., Mishig-Ochir, T., Lee, B.J.: Antimicrobial peptides: their physicochemical properties and therapeutic application. Arch Pharm Res 35(3), 409–413 (2012)

[43] Cock, P.J.A., Antao, T., Chang, J.T., Chapman, B.A., Cox, C.J., Dalke, A., Friedberg, I., Hamelryck, T., Kauff, F., Wilczynski, B., de Hoon, M.J.L.: Biopython: freely available Python tools for computational molecular biology and bioinformatics. Bioinformatics 25(11), 1422–1423 (2009). https://doi.org/10.1093/bioinformatics/btp163

[44] Pande, A., Patiyal, S., Lathwal, A., Arora, C., Kaur, D., Dhall, A., Mishra, G., Kaur, H., Sharma, N., Jain, S., et al.: Computing wide range of protein/peptide features from their sequence and structure. bioRxiv, 599126 (2019)

[45] Schaduangrat, N., Nantasenamat, C., Prachayasittikul, V., Shoombuatong, W.: Meta-iAVP: A Sequence-Based Meta-Predictor for Improving the Prediction of Antiviral Peptides Using Effective Feature Representation. Int J Mol Sci 20(22) (2019)

